# Mitofusin 2 controls mitochondrial and synaptic dynamics of suprachiasmatic VIP neurons and related circadian rhythms including sleep

**DOI:** 10.1101/2025.03.18.643991

**Authors:** Milan Stoiljkovic, Jae Eun Song, Hee-kyung Hong, Heiko Endle, Luis Varela, Jonatas Catarino, Xiao-Bing Gao, Zong-Wu Liu, Sabrina Diano, Jonathan Cedernaes, Joseph T. Bass, Tamas L. Horvath

## Abstract

Sustaining the strong rhythmic interactions between cellular adaptations and environmental cues has been posited as essential for preserving the physiological and behavioral alignment of an organism to the proper phase of the daily light/dark cycle. Here, we show that mitochondria and synaptic input organization of suprachiasmatic (SCN) vasoactive intestinal peptide (VIP)-expressing neurons show circadian rhythmicity. Perturbed mitochondrial dynamics achieved by conditional ablation of the fusogenic protein mitofusin 2 (Mfn2) in VIP neurons cause disrupted circadian oscillation in mitochondria and synapses in SCN VIP neurons leading to desynchronization of entrainment to the light/dark cycle in Mfn2 deficient mice that resulted in advanced phase angle of their locomotor activity onset, alterations in core body temperature and sleep-wake amount and architecture. Our data provide direct evidence of circadian SCN clock machinery dependence on high-performance Mfn2-regulated mitochondrial dynamics in VIP neurons for maintaining the coherence in daily biological rhythms of the mammalian organism.

## Introduction

The circadian rhythm is an endogenous oscillation in behavioral, physiological, and other bodily functions that have a periodicity of approximately 24 hours. The hypothalamic suprachiasmatic nucleus (SCN) is the master circadian pacemaker which by transcriptional/translational negative feedback loop-dependent gene expression and intrinsic neuronal circuitry entrains other brain regions and peripheral tissues to synchronize their activities in coherence with environmental time cues (1, 2). The SCN directly receives light input from the photosensitive retinal ganglion cells via the retinohypothalamic tract which heavily terminates into its ventral subdivision (SCN core) mainly containing vasoactive intestinal peptide (VIP)-expressing neurons (2). These neurons densely project outside the SCN core influencing other neuronal populations in its dorsal subdivision (SCN shell) that express VIP cognate receptors VPAC2. This anatomical feature reflects the vast influence VIP neurons have on the entire SCN circuit synchronization with the light/dark (LD) cycle and ultimately in sustaining circadian rhythms at the molecular, cellular, and behavioral levels (3–5). Accordingly, previous studies showed that developmental disruption of the VIP neurons or VPAC2 receptors can severely compromise both the synchrony in the SCN neuronal network and circadian amplitude (6–8), whereas targeted optogenetic activation of VIP neurons can adequately reset the circadian phase of the SCN ensemble (9).

The daily fluctuation of SCN gene expression and neuronal firing requires precise rhythmic regulation of local metabolic activity to meet its bioenergetic demand. Indeed, a rhythmic pattern in the expression levels of several genes involved in the control of metabolism was observed within SCN (10, 11). Furthermore, it was suggested that SCN metabolic rhythmicity is intertwined with mitochondrial morphological and functional plasticity (12–14). Ample evidence showed that in fulfilling cell-specific bioenergetic demand, mitochondria undergo tightly coordinated dynamic morphological changes reflected in their continuous fusion and fission (15, 16). Previous studies (17, 18) suggested the relationship between this mitochondrial adaptation and the circadian phase in the peripheral tissues, however, data directly connecting SCN clock machinery function with mitochondrial dynamics therein are scarce. Given the indispensable role of SCN VIP neurons in maintaining circadian rhythm (5), here we examine the impact of mitofusin 2 (Mfn2), a mitochondrial membrane protein involved in the control of the fusion process (19), on SCN VIP neuronal activity and downstream effects on behavior, thermoregulation, and sleep.

## Results

### Daily remodeling of synapses and mitochondria in the SCN VIP neurons is diminished in constant dark

To determine the impact of the SCN circadian oscillation in the LD cycle at the cellular level we first analyzed synapse density and mitochondrial morphology in SCN core VIP-expressing neurons of C57BL/6J mice at Zeitgeber times (ZT) 1, 7, 13, 19, (ZT0: light on; ZT12: light off). Our electron microscopy analysis revealed significantly higher number of synapses, and specifically excitatory synapses in SCN VIP neuronal somata at ZT7 compared to the other time points measured (Figure 1A). Analyzing mitochondria in SCN VIP neurons we found that they have a larger area and perimeter during the light phase, but their shape was more circular during the dark phase (Figure 1B-D, Supplemental Figure S1A and S1B). To better characterize changes in mitochondrial morphology, we established criteria based on measures of the perimeter (size) and aspect ratio (shape) of an individual mitochondrion. Then, we classified mitochondria in SCN VIP neurons as “big” and “small” groups and ”long” and “short” groups (Figure 1E) and found an inverse relationship between the “small-short” group and “big-long” group. The ratio of the “big-long” group increased during the light phase and decreased during the dark phase, while it has the opposite trend for the “small-short” group (Figure 1F). Mitochondrial density and coverage of the cytosolic area in SCN VIP neurons showed no change throughout the day (Supplemental Figure S1C-E).

**Figure 1.**
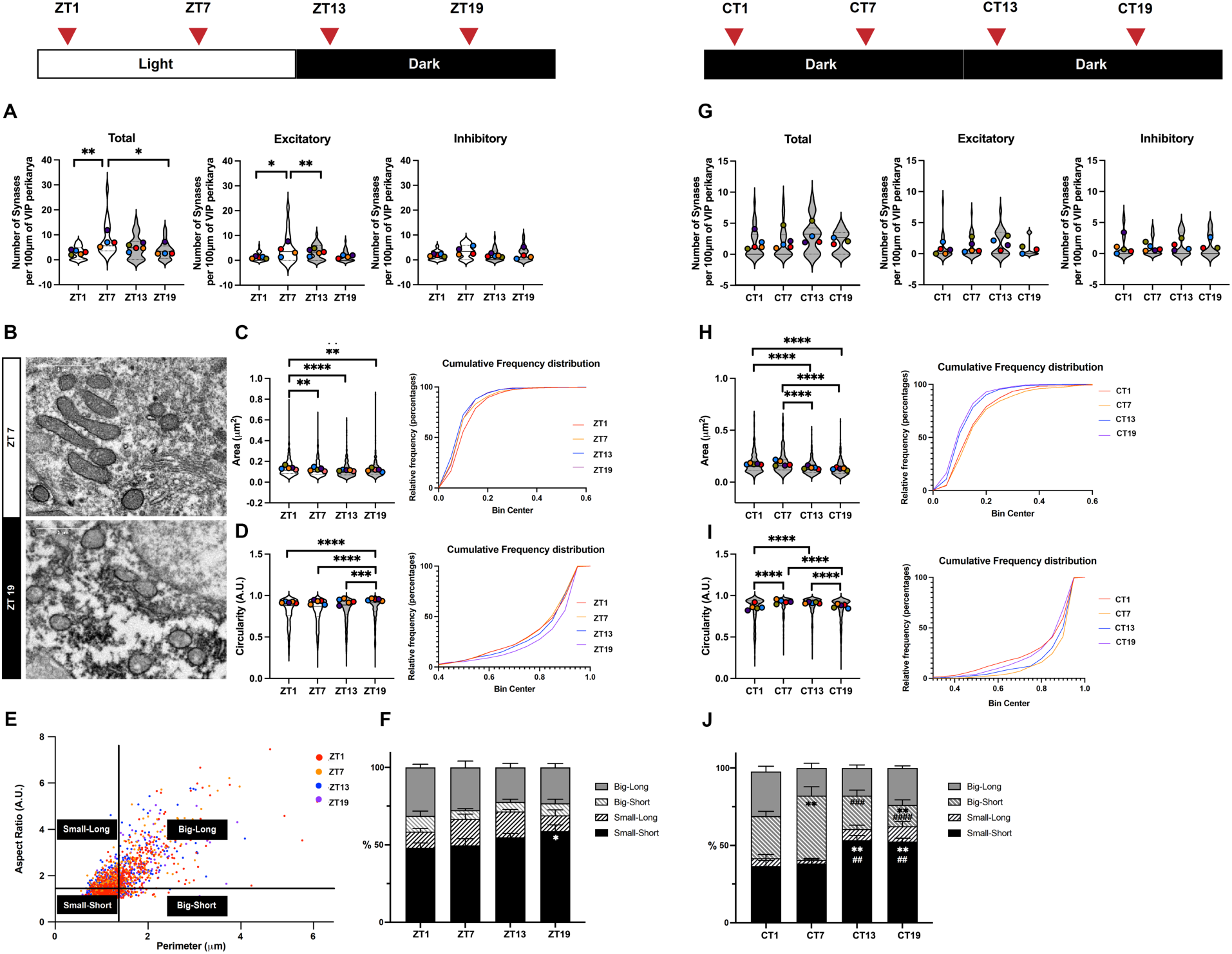
**Circadian rhythm of synaptic innervation and mitochondrial morphology of SCN VIP neurons in light/dark (LD) and constant darkness (DD) environment.** (**A-F**) Analyses of C57BL/6J mice housed in LD condition (ZT0: light on; ZT12: light off). (**A**) The density of synapses on SCN VIP neurons measured using electron microscope at ZT1, ZT7, ZT13, ZT19 under a normal light-dark (LD) cycle. (**B**) Representative electron microscopic images of mitochondria in SCN VIP neurons at ZT7 and ZT19. Scale bar = 1um. (**C**) Cross-sectional area and (**D**) circularity of mitochondria in SCN VIP neurons and their cumulative probability distributions at ZT1, ZT7, ZT13, ZT19 in LD. (**E**) Aspect ratio and perimeter of mitochondria in SCN VIP neurons. The average value of aspect ratio was used to determine the long/short group, and the average value of perimeter was used to determine the big/small group. (**F**) Percentage of four different groups of mitochondria in SCN VIP neurons at ZT1, ZT7, ZT13, ZT19 under LD conditions. Small/short group at ZT1 vs. ZT19; *P = 0.0372. (**G-J**) Analyses of C57BL/6J mice released into constant darkness (DD) for 48 hours. (**G**) The density of synapses on SCN VIP neurons measured using electron microscope at CT1, CT7, CT13, CT19 (DD condition). (**H**) Cross-sectional area and (**I**) circularity of mitochondria in SCN VIP neurons and their cumulative probability distributions at CT1, CT7, CT13, CT19 (DD condition). (**J**) Percentage of four different groups of mitochondria in SCN VIP neurons at CT1, CT7, CT13, CT19 (DD condition). Small/short group at CT1 vs. CT13, **P=0.013; CT1 vs. CT19, **P=0.015; CT7 vs. CT13, ^##^P=0.0069; CT7 vs. CT19, ^##^P=0.0086. Big/short group CT1 vs. CT7, **P=0.0072; CT1 vs. CT19, **P=0.0082; CT7 vs. CT13, ^###^P=0.0002; CT7 vs. CT19 ^####^P<0.0001. Approximately 5 cells per mice; 4-5 mice per time point. Supplemental Table S1 lists statistical information for each graph. *P<0.05; **P<0.01; ***P<0.005; ****P<0.0001.

Next, we repeated the experiment with the mice housed in constant darkness (DD) for 48 hours. The oscillation of synapse density in SCN VIP neurons was diminished with the absence of light signals (Figure 1G), suggesting that the remodeling of circadian synaptic innervation of these neurons is related to photic integration. During DD we observed an interesting change in the morphology of mitochondria. While the SCN VIP mitochondria size oscillations maintained the same pattern as in the LD condition, their shape drastically changed in the absence of light. The most noticeable change in mitochondrial morphology in the DD condition was the increase of “big-short” mitochondria overall, peaking at a circadian time point CT7 (corresponding to ZT7), in these neurons (Figure 1H-J, Supplemental Figure S1F and S1G). At circadian time points CT13 and CT19, the number of mitochondria was significantly higher but no change in the mitochondrial cytosolic coverage was found (Supplemental Figure S1H and S1I). These data indicate a shift of mitochondrial dynamics towards fission at ZT13 and ZT19 under constant darkness. While the “small-short” group maintained a similar circadian pattern, the “big-long” group changed its pattern in the DD condition when compared to the LD condition.

### Mfn2 deletion disrupts the daily remodeling of mitochondrial architecture in the SCN VIP neurons

Mitochondria undergo remodeling through fusion and fission to alter their distribution, morphology, and function in response to changes in their environment. To explore the effect of mitochondrial dynamics in the SCN circadian rhythmicity, we generated mice lacking mitofusin 2 (Mfn2^-/-^) in VIP neurons (Figure 2A). The loss of Mfn2 resulted in abnormal enlargement and circularity of mitochondria in SCN VIP neurons compared to the mitochondria from control mice in dark (ZT19) phase (Figure 2B and 2C). In the same phase, mitochondrial cytosolic coverage in Mfn2^-/-^ neurons did not change, but mitochondria density reduced likely due to an increase in their size (Figure 2D and 2E). These data indicate that the deletion of Mfn2 disrupts the mitochondrial dynamics in SCN VIP neurons. Unlike mitochondria in control SCN VIP neurons in which their perimeter and aspect ratio were positively correlated, a large portion of mitochondria in Mfn2^-/-^ neurons had their aspect ratio at 1 regardless of their perimeters (Figure 2F). Also, we found that SCN VIP neurons of Mfn2^-/-^ have increased circularity and perimeter during ZT19 (Figure 2G and 2H) suggesting a higher number of “big-short” mitochondria similar to that observed in mice housed under DD condition. In addition, no difference in mitochondria-ER contacts in SCN VIP neurons was observed between two groups of animals (Figure 2I).

**Figure 2.**
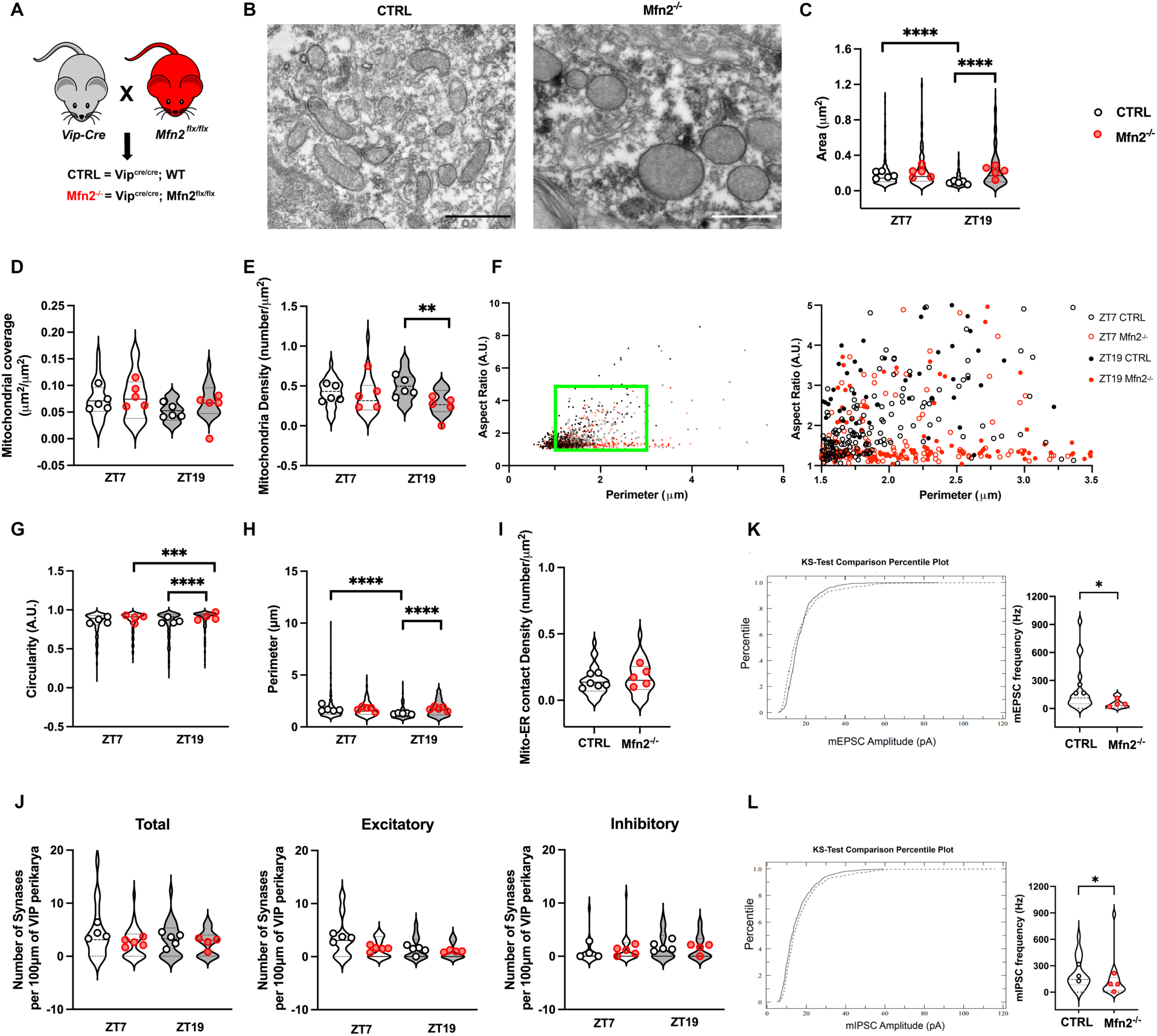
**Loss of Mfn2 alters mitochondrial morphology and synaptic innervations of SCN VIP neurons.** (**A**) Generation scheme for mice with Mfn-depleted SCN VIP neurons. Mfn2^flx/flx^ mice were crossed with VIP-Cre transgenic mice to generate VIP^cre/cre^;Mfn2^flx/flx^ (Mfn2^-/-^) and control VIP^cre/cre^ (CTRL) mice. (**B**) Representative electron microscopic images of mitochondria in VIP neurons in the SCN of CTRL and Mfn2^-/-^ at ZT7. (**C**) Cross-sectional area of mitochondria (**D**) mitochondrial coverage and (**E**) mitochondrial density of cytoplasm in SCN VIP neurons of CTRL and Mfn2 ^-/-^ mice at ZT7 and ZT19. (**F**) Aspect ratio and perimeter of mitochondria in SCN VIP neurons of CTRL and Mfn2^-/-^ mice. The green box indicates the location of zoomed-in version of the graph on the right. (**G**) circularity and (**H**) perimeter at ZT7 and ZT19, (**I**) mitochondria-ER contact per SCN VIP neuron of CTRL and Mfn2^-/-^ mice at ZT7, and (**J**) number of synapses in VIP cells at ZT7 and ZT19. Approximately 5 cells per mice; 4-6 mice per time point. (**K**) Cumulative distribution of miniature excitatory postsynaptic potentials (mEPSC) amplitude and frequency of SCN VIP neurons in CTRL (n = 20) and Mfn2^-/-^ (n = 16) mice measured at ZT7. (**L**) Cumulative distribution of miniature inhibitory postsynaptic potentials (mIPSC) amplitude and frequency of SCN VIP neurons in control (n = 19) and Mfn2^-/-^ (n = 20) mice measured at ZT7. Supplemental Table S1 lists statistical information for each graph. *P<0.05; **P<0.01; ***P<0.005; ****P<0.0001.

### Loss of Mfn2 in VIP neurons alters the synchronization of SCN neuronal activity

Along with mitochondria modifications in SCN VIP neurons of Mfn2^-/-^ mice, we analyzed daily rearrangements of the number of synapses in these neurons. In control animals, the total number of synapses, and particularly excitatory synapses, in SCN VIP neurons generally followed a similar pattern of reduction during the dark phase (ZT19) compared to the light phase (ZT7), consistent with trends previously observed in C57BL/6J mice, even though no significant differences were found. In contrast, Mfn2^-/-^ mice exhibited nearly the same number of SCN VIP synapses regardless of the diurnal phase (Figure 2J). This indicates impairment of circadian oscillation in SCN innervation in Mfn2^-/-^ mice likely because of deficient photic integration in SCN VIP neurons, which together with their “big-short” mitochondrial phenotype imply the involvement of mitochondrial dynamics in the light transduction process of SCN VIP neurons.

To further investigate the changes observed in Mfn2^-/-^ mice, miniature excitatory (mEPSC) and inhibitory (mIPSC) postsynaptic currents were recorded from SCN VIP neurons using brain slices. We detected a decrease in both mEPSC and mIPSC in Mfn2^-/-^ mice compared to controls (Figure 2K and 2L) at ZT7, indicating compromised synaptic functions of their VIP neurons. When measuring neuronal activation in the SCN VIP neurons at ZT7 and ZT19 by the analysis of cFos expression, we surprisingly observed higher VIP cFos positivity in Mfn2^-/-^ than in controls at ZT19 despite the findings of synaptic impairments in these animals (Figure 3A and 3C). Moreover, at the same time point we found the increased density of VPAC2 cFos expressing cells in the SCN nuclei of Mfn2^-/-^ mice (Figure 3B and 3D). This aligns with the previous observation (9) of marked cFos positivity induction in the entire SCN after optogenetic activation of VIP neurons. Overall, these findings suggest that increased SCN VIP activation in Mfn2^-/-^ mice is likely due to a compensatory mechanism potentially arising from Mfn2 loss-of-function during their developmental period. To clarify this further, future research employing conditional knockout models or tools for temporally controlled gene knockdown will be essential.

**Figure 3.**
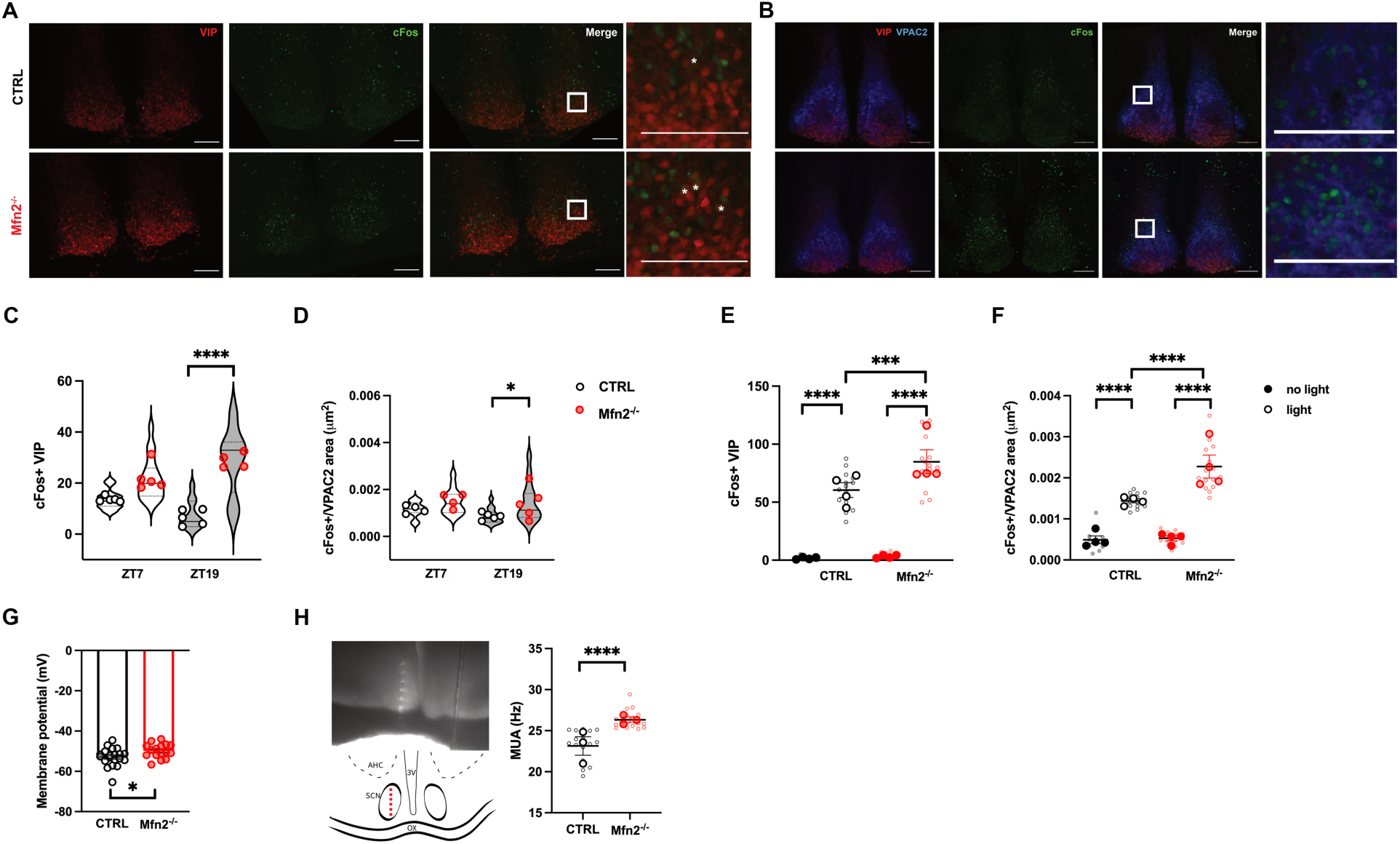
Loss of Mfn2 in VIP neurons alters the synchronization of SCN neuronal activity. (**A, C**) Representative confocal images of VIP neurons (red) and cFos (green) in the SCNs and number of cFos positive SCN VIP neurons of CTRL and Mfn2^-/-^ mice at ZT7 and ZT19. (**B, D**) Representative confocal images of VIP (red), VPAC2 (blue) and cFos (green) and density of cFos positive cells in the VPAC2-expressing area in the SCN of CTRL and Mfn2^-/-^ mice at ZT7 and ZT19. Analysis of 3 sections per mice (n = 4-5 mice/group). Scale bar = 100um. (**E**) Number of cFos expressing SCN VIP neurons and (**F**) the density of cFos expressing cells in VPAC2-expressing area in the SCN of CTRL and Mfn2^-/-^ after exposing to light for 1 hour at ZT13 (Light) or stayed in the dark (No light). Analysis of 4 sections per mice (n = 4-5 mice/group). (**G**) The membrane potential of SCN VIP neurons in CTRL (n = 20) and Mfn2^-/-^ (n = 16). (**H**) *Ex vivo* multiunit activity (MUA) recordings across the entire dorsoventral span of SCN in CTRL and Mfn2^-/-^ mice (n = 3/group). Microphotograph and diagram depict recording sites location in the SCN (labeling: 3V, third ventricle; OX, optic chiasm; AHC, anterior hypothalamic area central part). Supplemental Table S1 lists statistical information for each graph. *P<0.05; ***P<0.005; ****P<0.0001.

Next, we quantified the cFos expression in the SCN of Mfn2^-/-^ and control mice exposed to light at ZT13 for 1 hour. We observed increased light-induced activation of SCN VIP and VPAC2-expressing neurons in Mfn2^-/-^ (Figure 3E and 3F). To examine the underlying mechanism of this aberrant neuronal activation, we employed *ex vivo* electrophysiology recordings in two groups of animals to measure the membrane potential of the SCN VIP neurons and found that Mfn2^-/-^ mice have more depolarized membrane of VIP neurons than their control counterparts (Figure 3G). Also, using a multielectrode array placed across the whole dorsoventral dimension of SCN to record the neuronal spiking activity in the entire structure, we detected higher activity in Mfn2^-/-^ mice than in controls (Figure 3H). Although this result implies global dysfunction of their SCN neural circuitry, the role of VIP neurons is likely significant, as evidenced by the higher cFos expression and membrane depolarization. This may be attributed to the unique electrophysiological characteristics of VIP cells, which have a lower firing threshold and an increased spiking probability, as previously described (9, 20). Together, these data indicate that the functional neuronal changes caused by Mfn2 deletion in VIP neurons alter the synchronization in the SCN circuit and its response to light stimulation.

### Loss of Mfn2 in VIP neurons disrupts the onset of activity and alters diurnal rhythms

To determine the systemic consequences of Mfn2 deletion-induced alterations in SCN VIP synapses and mitochondria, we measured the circadian rhythm of locomotor activity under a normal LD, followed by constant darkness (DD). We found that both Mfn2^-/-^ and control mice have preserved circadian locomotor rhythmicity under DD, but Mfn2^-/-^ mice displayed an advanced phase angle of entrainment and shorter free-running period (Figure 4A and 4B). After a prolonged follow-up of wheel-running behavior in LD (Figure 4C), we observed altered activity onset in Mfn2^-/-^ mice relative to the dark phase. Compared to the control animals, they had shifted activity onset towards the light phase, resulting in higher activity counts during the light phase and statistically lower activity counts during the dark phase, but without changes in the total activity level (Figure 4D and 4E).

**Figure 4.**
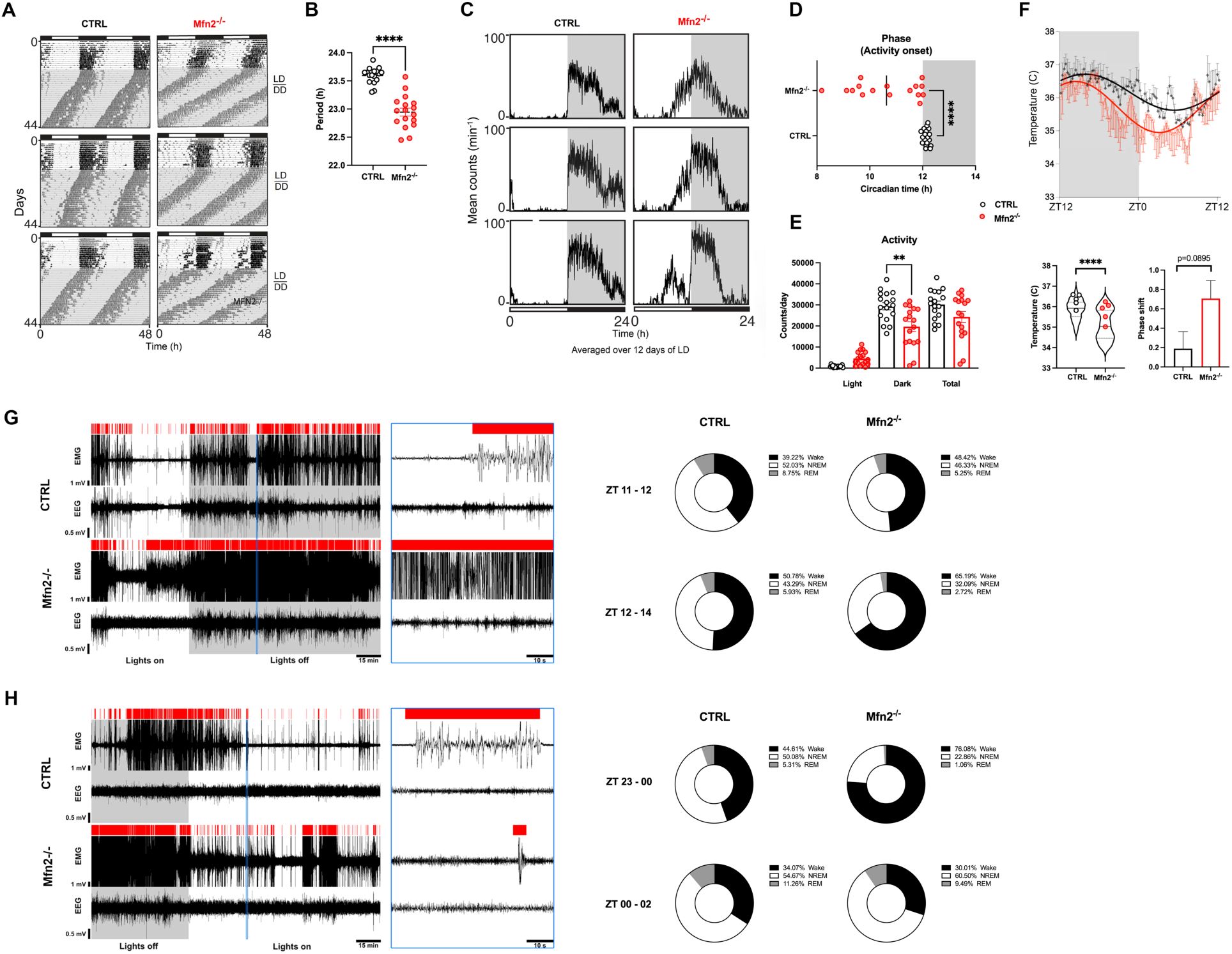
**Loss of Mfn2 in VIP neurons alters the diurnal activity rhythm and activity onset.** (**A**) Representative actogram showing wheel-running activity of CTRL and Mfn2^-/-^ mice. Mice were maintained on a 12:12 LD cycle in wheel cages for 16 days prior to release to DD. (**B**) Free-running endogenous circadian period distributions of CTRL and Mfn2^-/-^ mice. (**C**) Representative activity profile graph of wheel-running activity. Activity profile was averaged over 12 days of LD (days 5-16). (**D**) Phase of activity onset relative to the dark phase and (**E**) distribution of activity counts in the light or dark periods and total activity in CTRL and Mfn2^-/-^ mice averaged for days 5-16 in LD. (**F**) Core body temperature profiles over the entire LD cycle in the CRTL (black) and Mfn2^-/-^ mice (red). (**G-H**) Sleep-wake pattern measured during transitions from the light to dark phase (**G**) and the dark to light phase (**H**). Original EEG and EMG signals and distribution of automatically detected vigilant states in one CTRL and one Mfn2^-/-^ mouse during the last hour of light or dark phase, and the first two hours after shifting in the opposite phase. Panels at right show extracted signals (enclosed by blue lines) in 1 minute time interval. Red bars over the traces mark wake episodes. Pie charts show percentage distribution of wake (black), NREM (white) and REM (gray) episodes in CTRL and Mfn2^-/-^ mice (n = 5 for each). ZT23-00: CTRL vs Mfn2^-/-^, P=0.019 for wake; P=0.014 for NREM. Supplemental Table S1 lists statistical information for each graph. *P<0.05; **P<0.01; ****P<0.0001.

Next, when analyzing the core body temperature, we found a suggestively advanced phase shift with a decrease in average temperature values in Mfn2^-/-^ compared to control mice (Figure 4F). Further, assessing the sleep-wake pattern during the transition period from the light to dark phase revealed that these mice sleep 10–15% less on average, even though the significant difference between the two groups was not observed (Figure 4G). Conversely, the sleep-wake pattern during the transition from the dark to light phase was significantly disrupted in Mfn2^-/-^ mice, with a marked increase in wakefulness at the expense of NREM sleep during the final hour (ZT23–ZT0) of the dark phase (Figure 4H). Collectively, the results suggest asynchronous circadian entrainment of bodily biological rhythms to the environmental LD cycle in mice with impaired mitochondrial dynamics in SCN VIP cells due to Mfn2 deletion.

## Discussion

The distinctly timed output from SCN VIP neurons is essential for the circadian synchronization of daily biological rhythms (21). However, cellular mechanisms and factors which regulate the activity of these neurons with a high degree of fidelity necessary for maintaining the homeostatic rhythmicity of an organism have not been fully elucidated. Here, we analyze the cross-talk between mitochondria function in SCN VIP neurons and their circadian oscillatory output that governs biological rhythms of activity, body temperature and sleep. We first observed the association of the mitochondrial fusion and fission dynamics with photic integration in these neurons. When this adaptive structural arrangement of SCN VIP mitochondria is disrupted by Mfn2 downregulation, we detected a whole spectrum of synaptic morphological and electrophysiological alterations evoking desynchronization of the SCN circuitry. These changes consequently resulted in circadian behavior dysrhythmia in Mfn2^-/-^ mice with a significant impairment in their locomotor activity onset, core body temperature and sleep-wake regulation.

It has been recently reported that developmental disruption of the molecular clock within SCN VIP neurons, or their genetic ablation in adult mice led to profound changes in wheel-running activity, body temperature and sleep-wake rhythms (5). Interestingly, we showed that even less robust alteration in SCN VIP neurons, such as abnormality in their mitochondrial dynamics induced by Mfn2 loss-of-function is sufficient to trigger circadian dysrhythmia. This finding can be particularly important in the context of daily rhythmicity changes commonly observed during aging, and in age-related neurodegenerative diseases, given the evidence of declined mitochondrial functions therein (22, 23). These conditions are associated with impaired Mfn2-regulated mitochondrial fusion, which in turn affects the shape, distribution, and function of mitochondria as well as metabolic and bioenergetic properties of the cell (24, 25). In the SCN VIP neurons of Mfn2^-/-^ mice, we found abnormally enlarged, spherical-shaped mitochondria that lost their plasticity in daily morphological adaptations compared to that from controls. As a result, these Mfn2-deficient cells likely exhibit a reduced ability to sustain proper endogenous respiration and efficient energy production, as demonstrated in previous studies (26, 27). In turn, this can adversely impact the VIP/VPAC2 neuropeptidergic signaling axis within the SCN, leading to functional disruptions in its downstream effector targets and circadian regulation.

In accordance, in Mfn2^-/-^ mice, we found impaired circadian locomotor behavior characterized by an advanced shift in activity onset compared to control counterparts. This is consistent with the requisite role of SCN VIP neurons in regulating circadian locomotor activity given their direct projections to the dorsomedial hypothalamic nucleus which is shown to play an important role in this process (5, 28). Intriguingly, in contrast to the previously observed negative correlation between locomotor activity and SCN VIP activation (9), we did not detect any difference in total activity levels between mutant and control mice, even though increased activation of these neurons in Mfn2^-/-^ animals was found using cFos labeling. This possibly indicates that in the condition of compromised functioning of VIP neurons, the other SCN cell populations may be involved in circadian activity regulation in a compensatory manner.

Considering that the local SCN output is intimately integrated with hypothalamic centers involved in thermoregulation and sleep-arousal states (28–30), we also monitored the LD fluctuations of body temperature and sleep-wake pattern. In mice with Mfn2-depleted VIP neurons, we found defective core body temperature rhythm, similar to that observed after total ablation of these neurons in the SCN (5). Specifically, we noticed a daily average body temperature reduction in these mice compared to controls during the 24-hour LD cycle. This result can reflect increased inhibitory input from SCN VIP neurons of Mfn2^-/-^ animals, to the thermogenesis-promoting neurons in the dorsomedial hypothalamic nucleus-preoptic area circuit which is implicated in body temperature regulation based on ambient temperatures (31, 32). When testing whether disruption of mitochondrial dynamics in SCN VIP neurons affected sleep-wake patterns during LD and DL transition periods, we found that Mfn2^-/-^ mice exhibited significantly increased wakefulness during the last hour of the dark phase in the DL transition, compared to control mice. This likely result from increased activation of SCN VIP neurons that we observed in these animals during the dark phase, and conceptually aligns with previous findings of heightened total SCN activity during the waking episodes (33). Although measured in the restricted time window, i.e. only during phase transitions, this finding suggests altered sleep-wake behavior in Mfn2^-/-^ mice. The role of SCN VIP neurons in circadian sleep-wake regulation is ambiguous given the evidence showing that their activation does not directly modulate sleep and wake length and distribution across the day and night (5), as well as those reporting the essential role of their activation in sculpting nighttime but not daytime sleep-wake rhythm (29). Our findings regarding the involvement of these neurons in circadian gating of sleep and wake align with latter study, but given the impact of homeostatic regulatory factors it remains challenging to fully disentangle the active role of SCN VIP neurons in this process, underscoring the need for further investigation. In a broader view, these results on sleep-wake pattern change in Mfn2^-/-^ mice also provide new data for the emerging role of mitochondria in sleep control (34) which was not extensively studied thus far, in contrast to ample existing evidence showing how sleep impacts mitochondrial function and fitness (35).

Taken together our data indicate the importance of proper mitochondrial dynamics in SCN VIP neuronal output as a critical effector mechanism for shaping the biological rhythms of a mammalian organism in response to circadian daily cycle.

## Materials and Methods

### Sex as a biological variable

Our study examined male and female animals, and similar findings are reported for both sexes in all experiments.

### Mice

All mice were maintained in temperature and humidity-controlled rooms, in a 12-hr:12-hr light-dark cycle (light on at 7:00, light off at 19:00) unless otherwise stated. Food and water were provided ad libitum. Both VIP-IRES-Cre mice (Vip^tm1(cre)Zjh^//AreckJ) and their background control C57BL/6J mice were purchased from Jackson Laboratory. The VIP-IRES-Cre mice have Cre recombinase expression directed to *Vip*-expressing cells by the endogenous promoter/enhancer elements of the vasoactive intestinal polypeptide locus (*Vip*) on chromosome 10. To generate Mfn2^-/-^ mice they were crossed with Mitofusin 2 floxed mice (Mfn2^tm3Dcc^/Mmucd) available in our lab using the previously described approach (36, 37). To visualize VIP neurons, Cre-dependent tdTomato reporter control and Mfn2^-/-^ mice, aged 1.5-3 months, were used for histological analysis and *ex vivo* electrophysiological recordings. Mice between 2-6 months of age were used for *in vivo* electrophysiological recordings, body temperature, and wheel-running measurements .

### Immunostaining

Mice were anesthetized and transcardially perfused with ice-cold saline (0.9% NaCl) containing heparin (10mg/L) followed by a fixative solution (4% paraformaldehyde in 0.1M phosphate buffer (PB), pH 7.4). Brains were post-fixed overnight at 4°C. Vibratome 50um-thick sections were washed in PB for 15 min, then incubated in blocking solution (5% normal donkey serum in PB) with 0.3% Trition X-100 for 1 hour at room temperature. Sections were incubated with primary antibodies: rabbit anti-VPAC2 (1:300; ab183334, Abcam), goat anti-cFos (1:1000; sc-52-G, Santa Cruz) for 48 hours at 4°C, then washed 3 times in PB for 15 minutes each at room temperature and incubated with the respective secondary antibodies (donkey anti-rabbit 647, 1:700 (A31573, Thermo Fisher) and donkey anti-goat 488, 1:700 (A11055, Thermo Fisher)) overnight at 4°C. For VIP signal measurements, sections were incubated with rabbit anti-VIP antibody (1:500; 20077, Immunostar) for 72 hours at 4°C and then with donkey anti-rabbit 647 (1:500; A31573, Thermo Fisher) for 2 hours at room temperature. The following day, sections were washed and coverslipped using Vectashield mounting medium (H-1000, Vector Laboratories). All analyses were done using a Leica Stellaris 5WLL microscope (Leica Microsystems, Inc).

### Electron microscopy

Mice were anesthetized and first perfused with ice-cold saline (0.9% NaCl) containing heparin (10mg/L) followed by fixative solution (4% paraformaldehyde, 15% vol/vol picric acid, 0.1 vol/vol glutaraldehyde in 0.1M PB). Brains were post-fixed in a fixative solution without glutaraldehyde overnight at 4°C. For immunohistochemistry, 50 um-thick brain tissue sections that contained the SCN were first processed in 10% and 20% sucrose solution for 30 minutes each, and after a freeze-thaw step, they were incubated with rabbit anti-VIP antibody (1:1000; 20077, Immunostar) for 72 hours at 4°C. The next day, sections were washed, incubated with goat anti-rabbit HRP-conjugated antibody (Sigma 12-348; 1:200) for 1.5 hours, washed again, and developed with 3,3-diaminobenzidine (DAB). Subsequently, sections were osmicated and dehydrated in ethanol. Ultrathin sections cut using a Leica Ultra-Microtome were collected on Formvar-coated single-slot grids and analyzed with a Tecnai 12 Biotwin electron microscope (TEM-FEI).

### Image Analysis

Mitochondrial cross-sectional area, perimeter, circularity, and aspect ratio were calculated using the analyze particles function in ImageJ (https://imagej.net/imaging/particle-analysis).

Mitochondrial density was calculated by dividing the number of mitochondria in the cytosolic area. Mitochondrial coverage was calculated by dividing the total area of mitochondria by the cytosolic area. For synapse density, an investigator blinded to the experimental protocol measured the number of synapses, as previously described (38). For immunofluorescence image analyses, z-stack images of brain sections were collected to manually count the cFos expressing cells by an observer blinded to the experimental conditions. The measurement of the immunofluorescence intensity of VIP in SCNs was calculated using ImageJ. The background intensity of the SCN area was used to normalize the raw integrated density (RawIntDen) of the VIP antibody. The average value of two sides of SCNs was used per section.

### Electrophysiology

1. Sleep recording

Sleep recording was carried out in freely behaving male mice implanted with chronically indwelling electrodes. After achieving a surgical plane of isoflurane anesthesia (induction 4%, maintenance 1.5% in O_2_), mice were placed in Kopf stereotaxic frame, and stainless screw electrodes (Plastics One) were implanted under aseptic conditions over the right frontal and parietal cortices for electroencephalographic (EEG) recordings and wire electrode with suture pad inserted deep into the neck muscle tissue for monitoring electromyographic (EMG) activity. To serve as a reference two additional screws were placed over the cerebellum. All electrodes were joined to a miniature connector which was affixed to the skull using dental acrylic, and the skin incision was closed with nylon sutures. After surgery, each mouse was kept in a clean cage and all necessary postoperative care was taken. Following a 10-day recovery period, mice in their individual home cages were connected to slip-ring commutators with flexible cables and left undisturbed for habituation of at least 24 hours before EEG/EMG recording. All recordings were done during the transition periods in the light-dark cycle i.e., during the last hour of light or dark (ZT11-12 or ZT23-0) and the first two hours after shifting in the opposite phase (ZT12-14 or ZT0-2). The acquired EEG/EMG signal was filtered between 1 and 500 Hz using the A-M System (model 1800) with an additional notch filter at 60 Hz, simultaneously digitized at a rate of 1 kHz, and stored on a computer via CED Micro1401-3 interface and Spike2 version 9 software (Cambridge Electronic Design). Subsequent analyses were performed offline and blind to genotype in consecutive 5-second-long epochs using Spike2 scripts for automatic sleep staging and scoring according to standard criteria for rodents as described previously (39). Epochs scored as ambiguous (approx. <5%), occurring mainly due to occasional artifacts in the EEG signal, were excluded from the analysis.

2. Ex vivo SCN recording

Control and mutant Mfn2^-/-^ male mice were deeply anesthetized with isoflurane and decapitated at ZT7, their brains were rapidly removed, and coronal tissue sections of 300 µm thickness at the rostro-caudal level of SCN (40) were made using vibratome in an oxygenated (5% CO2 + 95% O2) cutting solution at 4 °C that contains (in mM): sucrose 220, KCl 2.5, CaCl2 1, MgCl2 6, NaH2PO4 1.25, NaHCO3 26, and glucose 10, pH 7.3 with NaOH. After preparation, slices were transferred to a recording chamber constantly perfused at a rate of 2 ml/min with artificial cerebrospinal fluid (containing in mM: NaCl 124, KCl 3, CaCl2 2, MgCl2 2, NaH2PO4 1.23, NaHCO3 26, glucose 10, pH 7.4 with NaOH) at 33 °C.

Whole-cell voltage-clamp (at −60 mV or 0 mV) was performed for measuring miniature excitatory and inhibitory postsynaptic currents (mEPSC and mIPSC) using Multiclamp 700 A Axon Instruments amplifier (Molecular Devices, LLC). The patch pipettes with a tip resistance of 4– 6 MΩ were made of borosilicate glass (World Precision Instruments) with a Sutter pipette puller (P-97) and filled with a pipette solution containing (in mM): K-gluconate 135, MgCl_2_ 2, HEPES 10, EGTA 1.1, Mg-ATP 2, Na_2_-phosphocreatine 10, and Na_2_-GTP 0.3, pH 7.3 with KOH. After the giga-ohm (GΩ) seal and whole-cell access were achieved, the series resistance (10–20 MΩ) was partially compensated by the amplifier. All data were sampled at 10 kHz, filtered at 3 kHz and analyzed with AxoGraph X event-detection package (AxoGraph, Inc), and plotted with Igor Pro software (WaveMetrics).

Multiunit activity (MUA) in SCN was recorded following slice preparation as above using a 16-channel silicone microelectrode linear array with 50 micrometers shank separation (A16×1 NeuroNexus) attached to the micromanipulator. The signals were amplified with an A-M System amplifier (model 3600), sampled at 10 kHz, and stored on the computer disc using the CED Micro1401-3 and Spike2. For every mouse, six recording sites spanning the entire dorsoventral dimension of SCN were identified from pictures taken at the end of each recording and subsequently analyzed with Spike2 software.

### Body temperature measurement

Mice were injected with buprenorphine (Ethiqa XR, 3.25 mg/kg, subcutaneously), anesthetized with isoflurane, abdominal skin was shaved, and a 1.5 cm incision was made at the abdominal midline. Real-time readable temperature telemetry transmitters (Anilogger system, Bodycap) were implanted into the intraperitoneal cavity ventral to the caudal arteries and veins but dorsal to the abdominal viscera. Incisions were sutured and treated by topical antibiotic, and each animal was placed in an individual cage and received subcutaneous injections of analgesic (Meloxicam 5 mg/kg) for two days. Following a 10-day recovery at an ambient temperature (22 ± 1 °C) and on a 12 h light 12 h dark cycle with food and water available *ad libitum*, their core body temperatures were measured every 15 minutes over the 24-hour period. Data were acquired using the portable telemetry receiver, transferred to a computer as an Excel file, and analyzed using Prism software.

### Wheel-running behavior

Mice were placed in individual wheel-running cages and allowed free access to food and water. Locomotor activity was recorded using the ClockLab Data Collection System (Actimetrics). Activity data was analyzed in 6-minute bouts using ClockLab software (Actimetrics). The free-running period was determined by line-fitting of activity onsets from data collected during the DD period.

### Statistical analyses

Statistical analyses were performed using unpaired two-sided t-tests, Mann-Whitney tests, one-way or two-way ANOVAs with Tukey’s post-hoc test, and Kruskal-Wallis tests with Dunn’s post-hoc test in GraphPad Prism 10 (GraphPad Software Inc). Normality was assessed using the Shapiro-Wilk test, and the choice of parametric or non-parametric tests was based on the results, as detailed in Supplemental Table S1. Cumulative distributions of mEPSC and mIPSC measurements were compared using the Kolmogorov-Smirnov test. For temperature variation analysis, a non-linear regression fit was applied using a sine function constrained to a 24-hour wavelength. A P value less than 0.05 was considered significant. The number of replicates for each experiment is specified in the figure legends.

### Study approval

All mouse studies were designed and performed in accordance with NIH guidelines and animal protocols approved by the Institutional Animal Care and Use Committees of both Yale and Northwestern Universities.

## Data availability

The data used to support the findings of this study are available within this article and Supplemental material file. Recordings of sleep, multiunit activity and wheel-running behavior have been deposited in the GIN open data management system [https://gin.g-node.org/Horvath_Lab/Mfn2-VIP-SCN.git].

## Author contributions

M.S., J.E.S., H-K.H., H.E., L.V., J.C., X-B.G., Z-W.L., S.D., J.C., J.T.B., and T.L.H conducted the experiments and analyzed data. M.S., J.E.S., and T.L.H designed the study and wrote the paper with input from all authors.

## Acknowledgments

This work was supported by NIH grants DK120891 to X-B.G., and T.L.H., DK126447, AG067329, AG051459 and AG082190 to T.L.H., and DK127800, DK132647, DK090625 and AG065988 to J.T.B.

## Supplemental material

Figures S1

Table S1

## References

1. Buhr ED, and Takahashi JS. Molecular components of the Mammalian circadian clock. Handb Exp Pharmacol. 2013(217):3–27.

2. Patton AP, and Hastings MH. The suprachiasmatic nucleus. Curr Biol. 2018;28(15):R816–R22.

3. Hastings MH, Maywood ES, and Brancaccio M. Generation of circadian rhythms in the suprachiasmatic nucleus. Nat Rev Neurosci. 2018;19(8):453–69.

4. Patton AP, Edwards MD, Smyllie NJ, Hamnett R, Chesham JE, Brancaccio M, et al. The VIP-VPAC2 neuropeptidergic axis is a cellular pacemaking hub of the suprachiasmatic nucleus circadian circuit. Nat Commun. 2020;11(1):3394.

5. Todd WD, Venner A, Anaclet C, Broadhurst RY, De Luca R, Bandaru SS, et al. Suprachiasmatic VIP neurons are required for normal circadian rhythmicity and comprised of molecularly distinct subpopulations. Nat Commun. 2020;11(1):4410.

6. Harmar AJ, Marston HM, Shen S, Spratt C, West KM, Sheward WJ, et al. The VPAC(2) receptor is essential for circadian function in the mouse suprachiasmatic nuclei. Cell. 2002;109(4):497–508.

7. Colwell CS, Michel S, Itri J, Rodriguez W, Tam J, Lelievre V, et al. Disrupted circadian rhythms in VIP- and PHI-deficient mice. Am J Physiol Regul Integr Comp Physiol. 2003;285(5):R939–49.

8. Hastings MH, Brancaccio M, and Maywood ES. Circadian pacemaking in cells and circuits of the suprachiasmatic nucleus. J Neuroendocrinol. 2014;26(1):2–10.

9. Mazuski C, Abel JH, Chen SP, Hermanstyne TO, Jones JR, Simon T, et al. Entrainment of Circadian Rhythms Depends on Firing Rates and Neuropeptide Release of VIP SCN Neurons. Neuron. 2018;99(3):555–63 e5.

10. Allen G, Rappe J, Earnest DJ, and Cassone VM. Oscillating on borrowed time: diffusible signals from immortalized suprachiasmatic nucleus cells regulate circadian rhythmicity in cultured fibroblasts. J Neurosci. 2001;21(20):7937–43.

11. Menger GJ, Lu K, Thomas T, Cassone VM, and Earnest DJ. Circadian profiling of the transcriptome in immortalized rat SCN cells. Physiol Genomics. 2005;21(3):370–81.

12. Isobe Y, Hida H, and Nishino H. Circadian rhythm of metabolic oscillation in suprachiasmatic nucleus depends on the mitochondrial oxidation state, reflected by cytochrome C oxidase and lactate dehydrogenase. J Neurosci Res. 2011;89(6):929–35.

13. Peek CB, Affinati AH, Ramsey KM, Kuo HY, Yu W, Sena LA, et al. Circadian clock NAD+ cycle drives mitochondrial oxidative metabolism in mice. Science. 2013;342(6158):1243417.

14. Sardon Puig L, Valera-Alberni M, Canto C, and Pillon NJ. Circadian Rhythms and Mitochondria: Connecting the Dots. Front Genet. 2018;9:452.

15. Liesa M, and Shirihai OS. Mitochondrial dynamics in the regulation of nutrient utilization and energy expenditure. Cell Metab. 2013;17(4):491–506.

16. Nasrallah CM, and Horvath TL. Mitochondrial dynamics in the central regulation of metabolism. Nat Rev Endocrinol. 2014;10(11):650–8.

17. Schmitt K, Grimm A, Dallmann R, Oettinghaus B, Restelli LM, Witzig M, et al. Circadian Control of DRP1 Activity Regulates Mitochondrial Dynamics and Bioenergetics. Cell Metab. 2018;27(3):657–66 e5.

18. Uchiyama Y. Circadian alterations in tubular structures on the outer mitochondrial membrane of rat hepatocytes. Cell Tissue Res. 1981;214(3):519–27.

19. Schrepfer E, and Scorrano L. Mitofusins, from Mitochondria to Metabolism. Mol Cell. 2016;61(5):683–94.

20. Obermayer J, Luchicchi A, Heistek TS, de Kloet SF, Terra H, Bruinsma B, et al. Prefrontal cortical ChAT-VIP interneurons provide local excitation by cholinergic synaptic transmission and control attention. Nat Commun. 2019;10(1):5280.

21. Paul S, Hanna L, Harding C, Hayter EA, Walmsley L, Bechtold DA, et al. Output from VIP cells of the mammalian central clock regulates daily physiological rhythms. Nat Commun. 2020;11(1):1453.

22. Lananna BV, and Musiek ES. The wrinkling of time: Aging, inflammation, oxidative stress, and the circadian clock in neurodegeneration. Neurobiol Dis. 2020;139:104832.

23. Sebastian D, Palacin M, and Zorzano A. Mitochondrial Dynamics: Coupling Mitochondrial Fitness with Healthy Aging. Trends Mol Med. 2017;23(3):201–15.

24. Chandhok G, Lazarou M, and Neumann B. Structure, function, and regulation of mitofusin-2 in health and disease. Biol Rev Camb Philos Soc. 2018;93(2):933–49.

25. Filadi R, Pendin D, and Pizzo P. Mitofusin 2: from functions to disease. Cell Death Dis. 2018;9(3):330.

26. Ding Y, Gao H, Zhao L, Wang X, and Zheng M. Mitofusin 2-deficiency suppresses cell proliferation through disturbance of autophagy. PLoS One. 2015;10(3):e0121328.

27. Yu SB, and Pekkurnaz G. Mechanisms Orchestrating Mitochondrial Dynamics for Energy Homeostasis. J Mol Biol. 2018;430(21):3922–41.

28. Chou TC, Scammell TE, Gooley JJ, Gaus SE, Saper CB, and Lu J. Critical role of dorsomedial hypothalamic nucleus in a wide range of behavioral circadian rhythms. J Neurosci. 2003;23(33):10691–702.

29. Collins B, Pierre-Ferrer S, Muheim C, Lukacsovich D, Cai Y, Spinnler A, et al. Circadian VIPergic Neurons of the Suprachiasmatic Nuclei Sculpt the Sleep-Wake Cycle. Neuron. 2020;108(3):486–99 e5.

30. Welsh DK, Takahashi JS, and Kay SA. Suprachiasmatic nucleus: cell autonomy and network properties. Annu Rev Physiol. 2010;72:551–77.

31. Tran LT, Park S, Kim SK, Lee JS, Kim KW, and Kwon O. Hypothalamic control of energy expenditure and thermogenesis. Exp Mol Med. 2022;54(4):358–69.

32. Uchida Y, Tokizawa K, and Nagashima K. Characteristics of activated neurons in the suprachiasmatic nucleus when mice become hypothermic during fasting and cold exposure. Neurosci Lett. 2014;579:177–82.

33. Deboer T, Vansteensel MJ, Detari L, and Meijer JH. Sleep states alter activity of suprachiasmatic nucleus neurons. Nat Neurosci. 2003;6(10):1086–90.

34. Sarnataro R, Velasco CD, Monaco N, Kempf A, and Miesenböck G. Mitochondrial origins of the pressure to sleep. *bioRxiv.* 2024:2024.02.23.581770.

35. Melhuish Beaupre LM, Brown GM, Braganza NA, Kennedy JL, and Goncalves VF. Mitochondria’s role in sleep: Novel insights from sleep deprivation and restriction studies. World J Biol Psychiatry. 2022;23(1):1–13.

36. Dietrich MO, Liu ZW, and Horvath TL. Mitochondrial dynamics controlled by mitofusins regulate Agrp neuronal activity and diet-induced obesity. Cell. 2013;155(1):188–99.

37. Stutz B, Nasrallah C, Nigro M, Curry D, Liu ZW, Gao XB, et al. Dopamine neuronal protection in the mouse Substantia nigra by GHSR is independent of electric activity. Mol Metab. 2019;24:120–38.

38. Varela L, Stutz B, Song JE, Kim JG, Liu ZW, Gao XB, et al. Hunger-promoting AgRP neurons trigger an astrocyte-mediated feed-forward autoactivation loop in mice. J Clin Invest. 2021;131(10).

39. Costa-Miserachs D, Portell-Cortes I, Torras-Garcia M, and Morgado-Bernal I. Automated sleep staging in rat with a standard spreadsheet. J Neurosci Methods. 2003;130(1):93–101.

40. Paxinos G, and Franklin KBJ. Paxinos and Franklin’s the Mouse Brain in Stereotaxic Coordinates. Cambridge, Massachusetts, United States: Academic Press; 2019.

