## Supplemental material for "Mitofusin 2 controls mitochondrial and synaptic dynamics of suprachiasmatic VIP neurons and related circadian rhythms including sleep"

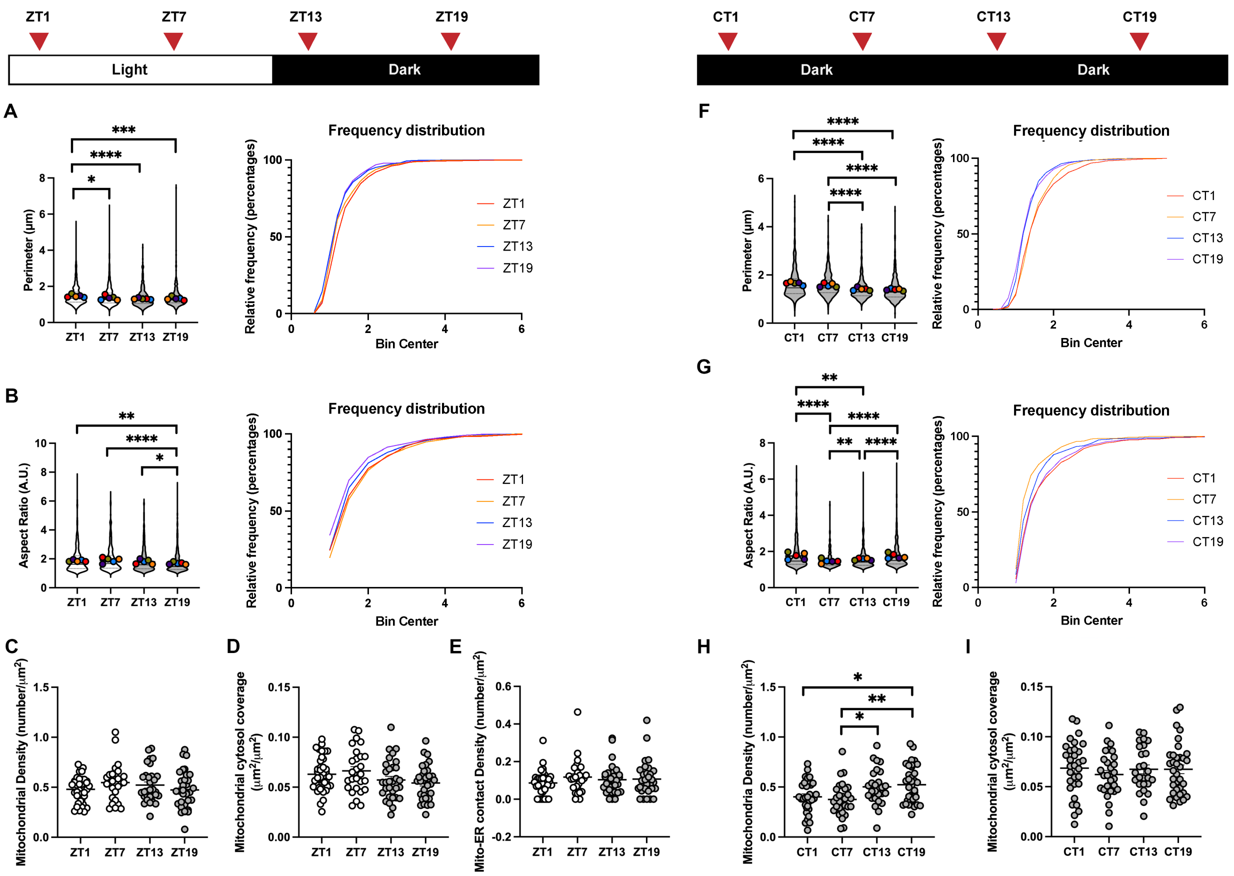


### Figure S1. Mitochondrial morphology of SCN VIP neurons in light-dark (LD) and constant darkness (DD) environment.

(**A-E**) Electron microscopy image analyses of SCN VIP neurons from C57BL/6J mice housed in LD condition (ZT0: light on; ZT12: light off). (**A**), Cross sectional perimeter; (**B**), Aspect ratio of mitochondria in SCN VIP neurons and their cumulative probability distributions at ZT1, ZT7, ZT13, ZT19 in LD. (**C**), Mitochondrial density; (**D**), Mitochondrial cytosol coverage; (**E**), Mitochondria-ER contact per SCN VIP neuron at ZT1, ZT7, ZT13, ZT19 in LD. (**F-I**) Electron microscopy image analyses of SCN VIP neurons from C57BL/6J mice housed in DD condition for 48 hours. (**F**), Cross sectional perimeter; (**G**), Aspect ratio of mitochondria in SCN VIP neurons and their cumulative probability distributions at CT1, CT7, CT13, CT19 in DD. (**H**), Mitochondrial density; (**I**), Mitochondrial cytosol coverage per SCN VIP neuron at CT1, CT7, CT13, CT19 in DD. Approximately 5 cells per mice; 5 mice per time point. Supplemental Table S1 lists statistical information for each graph. $^{*}$P<0.05; $^{**}$P<0.01; $^{***}$P<0.005; $^{****}$P<0.0001.

**Table S1. Statistics underlying Figures and Supplemental Figure**

| Figure | Data structure | Type of test | Test result |
|---|---|---|---|
| 1A total | heterogenous distribution | Kruskal-Wallis with Dunn's test | H = 14.17, P=0.003 |
| 1A excit | heterogenous distribution | Kruskal-Wallis with Dunn's test | H = 15.90, P=0.001 |
| 1A inhib | heterogenous distribution | Kruskal-Wallis with Dunn's test | H = 6.83, P=0.078 |
| 1C | heterogenous distribution | Kruskal-Wallis with Dunn's test | H = 39.24, P<0.001 |
| 1D | heterogenous distribution | Kruskal-Wallis with Dunn's test | H = 30.51, P<0.001 |
| 1F | normal distribution | 2-way ANOVA with Tukey's test | $F_{(15, 60)}$ = 55.10, P<0.001 |
| 1G total | heterogenous distribution | Kruskal-Wallis with Dunn's test | H = 3.59, P=0.309 |
| 1G excit | heterogenous distribution | Kruskal-Wallis with Dunn's test | H = 5.06, P=0.167 |
| 1G inhib | heterogenous distribution | Kruskal-Wallis with Dunn's test | H = 1.24, P=0.743 |
| 1H | heterogenous distribution | Kruskal-Wallis with Dunn's test | H = 129.5, P<0.001 |
| 1I | heterogenous distribution | Kruskal-Wallis with Dunn's test | H = 89.59, P<0.001 |
| 1J | normal distribution | 2-way ANOVA with Tukey's test | $F_{(15, 72)}$ = 27.85, P<0.001 |
| 2C | heterogenous distribution | Kruskal-Wallis with Dunn's test | H = 172.7, P<0.001 |
| 2D | normal distribution | 1-way ANOVA with Tukey's test | $F_{(3, 101)}$ = 3.10, P=0.03 |
| 2E | heterogenous distribution | Kruskal-Wallis with Dunn's test | H = 14.07, P=0.002 |
| 2G | heterogenous distribution | Kruskal-Wallis with Dunn's test | H = 44.88, P<0.001 |
| 2H | heterogenous distribution | Kruskal-Wallis with Dunn's test | H = 131.7, P<0.001 |
| 2I | heterogenous distribution | Mann Whitney test | U = 394, P=0.311 |
| 2J total | heterogenous distribution | Kruskal-Wallis with Dunn's test | H = 1.002, P=0.801 |
| 2J excit | heterogenous distribution | Kruskal-Wallis with Dunn's test | H = 7.47, P=0.058 |
| 2J inhib | heterogenous distribution | Kruskal-Wallis with Dunn's test | H = 3.87, P=0.275 |
| 2K amp | heterogenous distribution | Kolmogorov-Smirnov test | D = 0.091, P=0.014 |
| 2K freq | heterogenous distribution | Mann Whitney test | U = 80, P=0.01 |
| 2L amp | heterogenous distribution | Kolmogorov-Smirnov test | D = 0.160, P<0.001 |
| 2L freq | heterogenous distribution | Mann Whitney test | U = 118, P=0.043 |
| 3C | heterogenous distribution | Kruskal-Wallis with Dunn's test | H = 25.89, P<0.001 |
| 3D | normal distribution | 1-way ANOVA with Tukey's test | $F_{(3, 51)}$ = 3.86, P=0.014 |
| 3E | normal distribution | 1-way ANOVA with Tukey's test | $F_{(3, 53)}$ = 126.2, P<0.001 |
| 3F | normal distribution | 1-way ANOVA with Tukey's test | $F_{(3, 55)}$ = 95.08, P<0.001 |
| 3G | normal distribution | unpaired t test | t = 2.23, df=34, P=0.032 |
| 3H | heterogenous distribution | Mann Whitney test | U = 0, P<0.001 |
| 4B | normal distribution | unpaired t test | t = 8.20, df=32, P<0.001 |
| 4D | normal distribution | unpaired t test | t = 5.08, df=32, P<0.001 |
| 4E | normal distribution | 1-way ANOVA with Tukey's test | $F_{(5, 96)}$ = 51.48, P<0.001 |
| 4F temp | heterogenous distribution | Mann Whitney test | U = 2256, P<0.001 |
| 4F phase | normal distribution | unpaired t test | t = 2.02, P=0.089 |
| 4G | normal distribution | multiple unpaired t tests with Welch corrections | ZT11-12:P=0.22(W); P=0.47(NR);P=0.39(R); ZT12-14:P=0.21(W); P=0.25(NR);P=0.18(R) |
| 4H | normal distribution | multiple unpaired t tests with Welch corrections | ZT23-0:P=0.02(W);P=0.014(NR);P=0.154(R); ZT0-2:P=0.656(W); P=0.553(NR);P=0.449(R) |
| S1A | heterogenous distribution | Kruskal-Wallis with Dunn's test | H = 28.62, P<0.001 |
| S1B | heterogenous distribution | Kruskal-Wallis with Dunn's test | H = 25.21, P<0.001 |
| S1C | heterogenous distribution | Kruskal-Wallis with Dunn's test | H = 3.29, P=0.349 |
| S1D | normal distribution | 1-way ANOVA with Tukey's test | $F_{(3, 125)}$ = 2.55, P=0.005 |
| S1E | heterogenous distribution | Kruskal-Wallis with Dunn's test | H = 2.05, P=0.561 |
| S1F | heterogenous distribution | Kruskal-Wallis with Dunn's test | H = 94.29, P<0.001 |
| S1G | heterogenous distribution | Kruskal-Wallis with Dunn's test | H = 73.02, P<0.001 |
| S1H | normal distribution | 1-way ANOVA with Tukey's test | H = 1.28, P=0.732 |
| S1I | heterogenous distribution | Kruskal-Wallis with Dunn's test | $F_{(3, 117)}$ = 5.39, P=0.001 |

Shapiro-Wilk test was used for assessing data normality and lognormality.
